# Optimising colour for camouflage and visibility using deep learning: the effects of the environment and the observer’s visual system

**DOI:** 10.1101/428193

**Authors:** J.G. Fennell, L. Talas, R.J. Baddeley, I.C. Cuthill, N.E. Scott-Samuel

**Affiliations:** School of Psychological Science, University of Bristol, 12a Priory Road, Bristol, BS8 1TU, UK; School of Biological Sciences, University of Bristol, Bristol Life Sciences Building, 24 Tyndall Avenue, Bristol, BS8 1TQ

**Author notes:** Corresponding author (JGF). JGF - Data Curation, Formal analysis, Investigation, Project Administration, Software, Validation, Visualisation, Writing – original draft, Writing – review & editing. LT - Data Curation, Formal analysis, Investigation, Software, Validation, Visualisation, Writing – review & editing. RJB, ICC and NESS - Conceptualisation, Funding Acquisition, Writing – review & editing.

## Abstract

Avoiding detection can provide significant survival advantages for prey, predators, or the military; conversely, maximising visibility would be useful for signalling. One simple determinant of detectability is an animal’s colour relative to its environment. But identifying the optimal colour to minimise (or maximise) detectability in a given natural environment is complex, partly because of the nature of the perceptual space. Here for the first time, using image processing techniques to embed targets into realistic environments together with psychophysics to estimate detectability and deep neural networks to interpolate between sampled colours, we propose a method to identify the optimal colour that either minimises or maximises visibility. We apply our approach in two natural environments (temperate forest and semi-arid desert) and show how a comparatively small number of samples can be used to predict robustly the most and least effective colours for camouflage. To illustrate how our approach can be generalised to other non-human visual systems, we also identify the optimum colours for concealment and visibility when viewed by simulated red-green colour-blind dichromats, typical for non-human mammals. Contrasting the results from these visual systems sheds light on why some predators seem, at least to humans, to have colouring that would appear detrimental to ambush hunting. We found that for simulated dichromatic observers, colour strongly affected detection time for both environments. In contrast, trichromatic observers were more effective at breaking camouflage.

**Author Summary:** Being the right colour is important in a natural and built environment, both for hiding (and staying alive) or being seen (and keeping safe). However, empirically establishing what these colours might be for a given environment is non-trivial, depending on factors such as size, viewing distance, lighting and occlusion. Indeed, even with a small number of factors, such as colour and occlusion, this is impractical. Using artificial intelligence techniques, we propose a method that uses a modest number of samples to predict robustly the most and least effective colours for camouflage. Our method generalises for classes of observer other than humans with normal (trichromatic) vision, which we show by identifying the optimum colours for simulated red-green colour-blind observers, typical for non-human mammals, as well as for different environments, using temperate forest and semi-arid desert. Our results reveal that colour strongly affects detection time for simulated red-green colour-blind observers in both environments, but normal trichromatic observers were far more effective at breaking camouflage and detecting targets, with effects of colour being much smaller. Our method will be an invaluable tool, particularly for biologists, for rapidly developing and testing optimal colours for concealment or conspicuity, in multiple environments, for multiple classes of observer.

## Introduction

Recently, interest in camouflage among evolutionary biologists has grown considerably [1], and many of the basic principles of how to conceal oneself have become far clearer. The range of research is wide and empirical support has been provided for many of the diverse strategies employed in the animal kingdom. Studies measuring the effectiveness of camouflage tend follow the same basic format: a small number of colours, patterns or colour/pattern combinations are generated that capture the proposed camouflage principles; and then the utility of the camouflage is evaluated, perhaps in the field by measuring predation rates, or in the laboratory measuring detection speed and accuracy, identification ability, or capture rate, using either human or non-human subjects. The same basic method is also used in the assessment of military camouflage [e.g. 2-5]. If the goal is to compare only a few colours/patterns in a given context then this strategy has much to commend it, being both simple to analyse and easy to understand. However, if the question is “what is the optimal camouflage strategy to employ in a given context?”, the approach is ineffective: the range of possible patterns is too large.

Optimal camouflage depends on a diverse range of factors: size, viewing distance, height above the ground, lighting, occlusion, the nature and variability of the environment, as well as the characteristics of the visual system of the observer [1,6-10]. The optimal colours and patterns may also vary depending on the mechanism by which the camouflage acts, whether to hinder detection, identification, selection or capture [1,10]. Consequently, in the animal kingdom, the range of camouflage patterns and strategies is wide; in human applications (e.g. military), the range of potential patterns is even wider because pattern generation is not constrained by biological mechanisms [11]. A reliable, systematic means of finding the optimal colouration and pattern for either minimising (camouflage) or maximising (conspicuity) visibility for a given range of environments would have wide applicability.

Here we concentrate on one simple but important characteristic of camouflage: its colour. Partly this is because colour is obviously an important property in determining the visibility of a target but, more importantly, because the space of all possible colours is far larger than traditionally explored. If colour could be characterised by a single dimension (say simply its luminance), then identifying the most (or least) concealing colours in a given context would be straightforward. We could systematically vary the colour along this single dimension and use a principled method to assess its visibility: finding maxima (and minima) in one dimensional spaces is simple. In contrast, even though colour is relatively low dimensional, an exhaustive evaluation of all colours, even if done at a coarse scale, scales badly: the number of locations required in colour space increases exponentially with dimensionality. We would like a method that could scale to this number of dimensions, and if this works, hopefully scale it to additional dimensions (such as texture).

Equally, investigating only those colours and patterns seen in nature [e.g. 12] omits possibilities that became extinct without leaving a fossil record, or those that evolution has not realised because of phylogenetic or developmental constraints. To address the challenges presented by a large multi-dimensional parameter space, without needing to impose artificial constraints, we propose a method for identifying optimal colours, patterns or colour/pattern combinations in any given context that uses deep neural networks. Specifically, we use them to interpolate smoothly between the colours that have been (noisily) tested, to other colours that have not.

Neural networks have been used for finding structure in unlabelled data (unsupervised learning) [13]; classification of inputs based on previously labelled data [14]; or regression (predicting real valued measurements) [15]. In some ways this may be considered a “sledge-hammer” approach: usually the simple problems such as interpolating a three-dimensional space would be dealt with a simpler method such as Gaussian process-based smoothing [16]. However, such methods often inherit strong assumptions, such as constant variability/noise for all values of the function. While these assumptions may be reasonable for the simple three-dimensional problems studied here, deep neural networks are a more general solution and can potentially be applied to more complicated spaces. Here we use deep neural networks to implement non-linear regression and use it, after training, to interpolate between measured inputs and predict responses for unseen inputs.

While the data sampling requirements of our method are modest compared with contemporary ‘big data’ standards, they are nevertheless large enough to preclude field trials. Using computer presentation and human participants, we can change stimuli rapidly and accurately capture the reaction times taken to identify them. However, because many camouflage strategies (such as concealment of shape based on countershading [17]) simply do not make sense in a uniformly illuminated two-dimensional world, and many objects are effectively impossible to conceal unless partly hidden by the foreground, we built our stimuli using multiple layers in order to achieve some level of realism. Our stimuli superimposed three layers: 1) a foreground occlusion layer; 2) a target layer; and 3) a background layer. We then used these stimuli to construct a visual search task that in some sense matches the task of predators and prey, albeit without active movements through the environment. In this way we can control each of the dimensions of interest.

To provide some confidence that the approach generalises, we demonstrate our method using human participants to identify targets of single colour in trichromat and simulated dichromat conditions in two natural environments. Dichromatic colour is straightforward to simulate for trichromats, using image processing; though the downside is the lack of the lifetime’s experience of dichromacy that a natural protanope has, something returned to in the discussion. The natural environments we used were temperate forest and semi-arid desert. We show: (i) that our methods allow rapid presentation of coloured objects embedded in realistic environments; (ii) how neural networks can be combined with bootstrap techniques to provide a statistical characterisation of the visibility function (the mapping between the colour of an object and its geometric mean detection time); and (iii) that the optimally concealed and conspicuous targets depend not only on the environment they are embedded in, but also on the nature of the visual system of the observer.

## Results

### Training networks

Human reaction time data for each condition were combined to provide a trichromat dataset and dichromat dataset (each one consisting of 500 trials x 10 participants = 5000) for each geographical location. In order to be able to interpolate and predict reaction times for target colours that had not been sampled during the experiment, and to take account of inter-subject variability in responses, residual deep neural network models were built using the high-level neural network API Keras 2.1.2 [18] running on top of neural network library TensorFlow 1.5.0 [19], separately for each combination of geographic location and chromatic condition. Inputs to the networks were the colour of the target (as RGB triplets), occlusion level and an one-hot array for participant IDs, while the output was the predicted reaction time. An alternative colour space such as CIELab or HSV could have been used, however, as neural networks form their own internal representations of distances [20] the choice of colour space is irrelevant. To provide for a measure of accuracy in our predictions (an estimate of standard error) we created 100 bootstraps of our networks. The bootstrap method is a test or metric that uses random sampling with replacement. The Bootstrap allows assignment of accuracy measures, defined here in terms of variance and is particularly useful when the value of interest is, as in the present case, a complicated function [21]. By averaging the bootstrapped networks predictions we calculate both a data dependent smoothing of the reaction time function and an estimate of our certainty of its estimate. Each network was trained on a random sample of 90% of the data and validated with the remaining 10%. In order to establish the number of residual blocks to use, networks were trained with two, four and six residual blocks. By comparing validation loss we found that four residual blocks gave the lowest error in all conditions, and accordingly this configuration was used. Further information for details of network configuration (together with how residual blocks are defined), comparing validation losses and parameters can be found in Supplementary information.

### Network predictions

The training process resulted in 100 models for each geographical location and chromatic condition, i.e. four sets of 100 models. Using each set of trichromat bootstrapped models, we submitted 16,777,216 colour samples (the whole RGB gamut), collecting predicted reaction times for each. Similarly, for each set of dichromat bootstrapped models, we submitted the entire simulated dichromat gamut (64,229 colour samples). All networks predictions were made using 37.5% occlusion, which was the average occlusion level across experimental trials. From the bootstrap values, we found the easiest and hardest to see colours by averaging across the reaction times for the 100 bootstraps per condition. In order to compare predicted reaction times for colours within and between geographical locations, statistics were calculated using random permutation tests, all based on 100,000 resamples. The predicted detection times were significantly lower in the dichromat treatment than in the trichromat treatment for both locations (all *p* < .0001, Figure 1). The difference between the hardest and easiest to find colours within geographical locations was also found to be significant for both chromatic conditions (all *p* < .0001). P-values were adjusted for multiple comparisons with False Discovery Rate [22].

**Figure 1.**
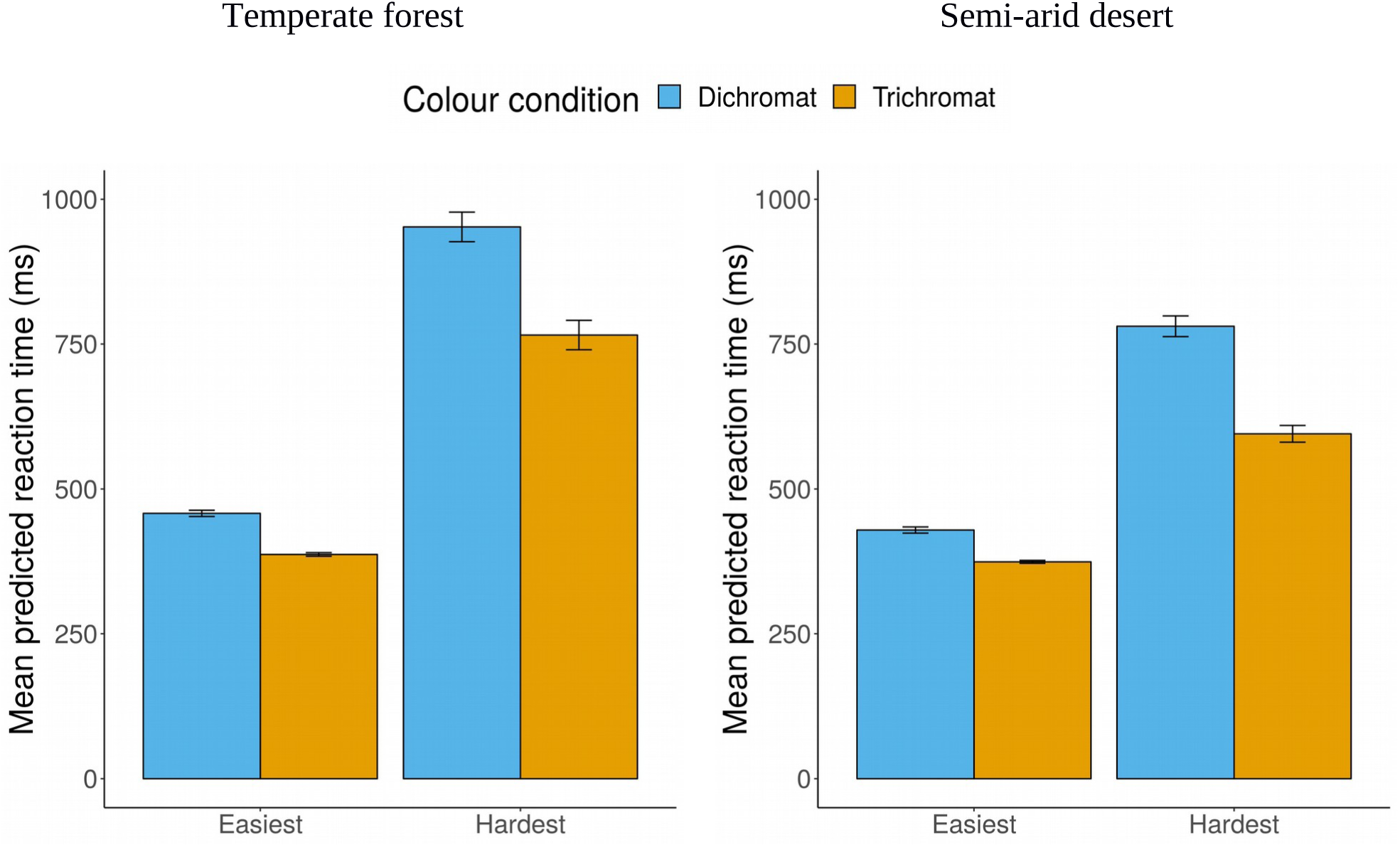
Mean predicted reaction times from bootstrapped neural networks. Error bars represent 1 SEM.

To provide an illustration of how networks predict reaction times to the respective colour gamuts, we created polar plots showing the predicted reaction times with respect to the hardest to see colour (Figure 2). For trichromats in the forest environment, the top left panel of Figure 2 shows that a shade of dark green/khaki is the hardest to find and shades of red, magenta and neon green are the easiest to find. In the desert trichromat setting (bottom left panel) trichromats find shades of beige (plainly reminiscent of light and dark sand) the most difficult and again neon green the easiest to find. For the forest dichromat condition, top right panel of Figure 2 clearly shows that a dark olive shade is the hardest to find, while blue, white and bright yellow stand out most. In the desert trichromat condition (bottom right panel) the hardest to find colour is a lighter beige shade and the easiest light blue. The white spaces containing no colour points in Figure 2 illustrate that no (or few) points were found at those reaction times. In other words, using the top left panel for forest trichromat, the white space at around 220° indicates that none of the light blue hues were difficult to find.

**Figure 2.**
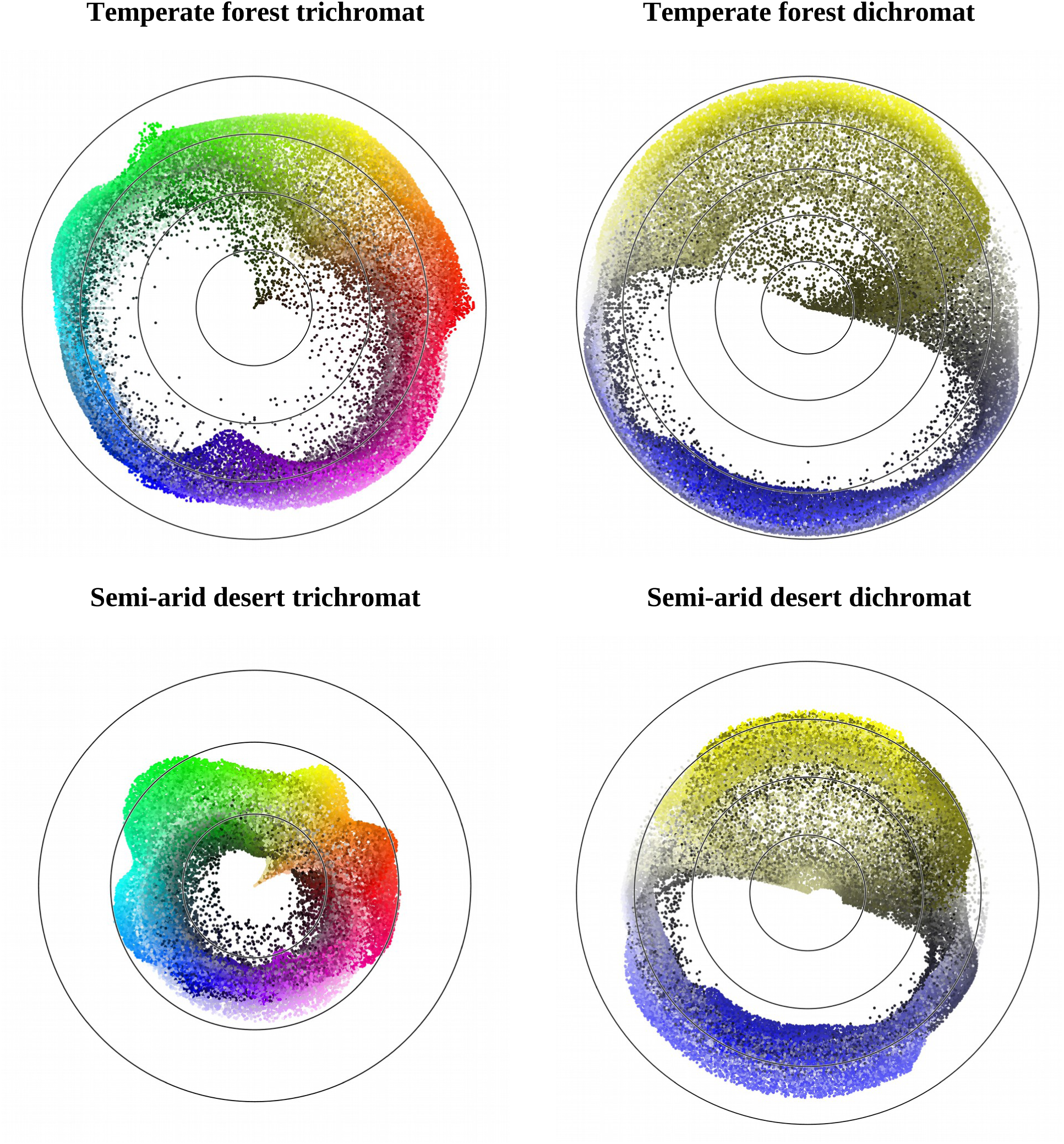
Polar plots showing the predicted reaction time difference from the hardest to find colours in the centre of the plot for each geographical location and chromatic condition. The angle is given by hue, representing red starting at 0°, yellow (from 60°), green (120°), cyan (180°), blue (240°) and magenta (300°). Distance is given by the difference in reaction time from the hardest to find colour. From the hardest to see colour in the centre of the plot, each contour represents an additional 100ms predicted reaction time.

### Validation

To provide a level of confirmation that dichromat colours are harder to see than trichomat colours and that the ordinality (in terms of reaction times) was comparable, we conducted a simple validation experiment using the same method as the original data collection. The only difference in the procedure was that colour choices were confined to three categories: easiest, intermediate (chosen halfway between reaction time extremes) and hardest. Twenty-five colours were chosen arbitrarily within a 25 ms margin of the predicted reaction times associated with each category (Figure 3 illustrates the stimuli used). This was done because, although we have identified a single hardest and easiest colour for each condition, characterising an entire function simply by its maxima and minima fails to capture the function completely. Data were analysed using Generalised Linear Mixed Models and we found that the results of the validation were consistent with predictions from the neural networks. Further information for details of the data analysis can be found in Supplementary information.

**Figure 3.**
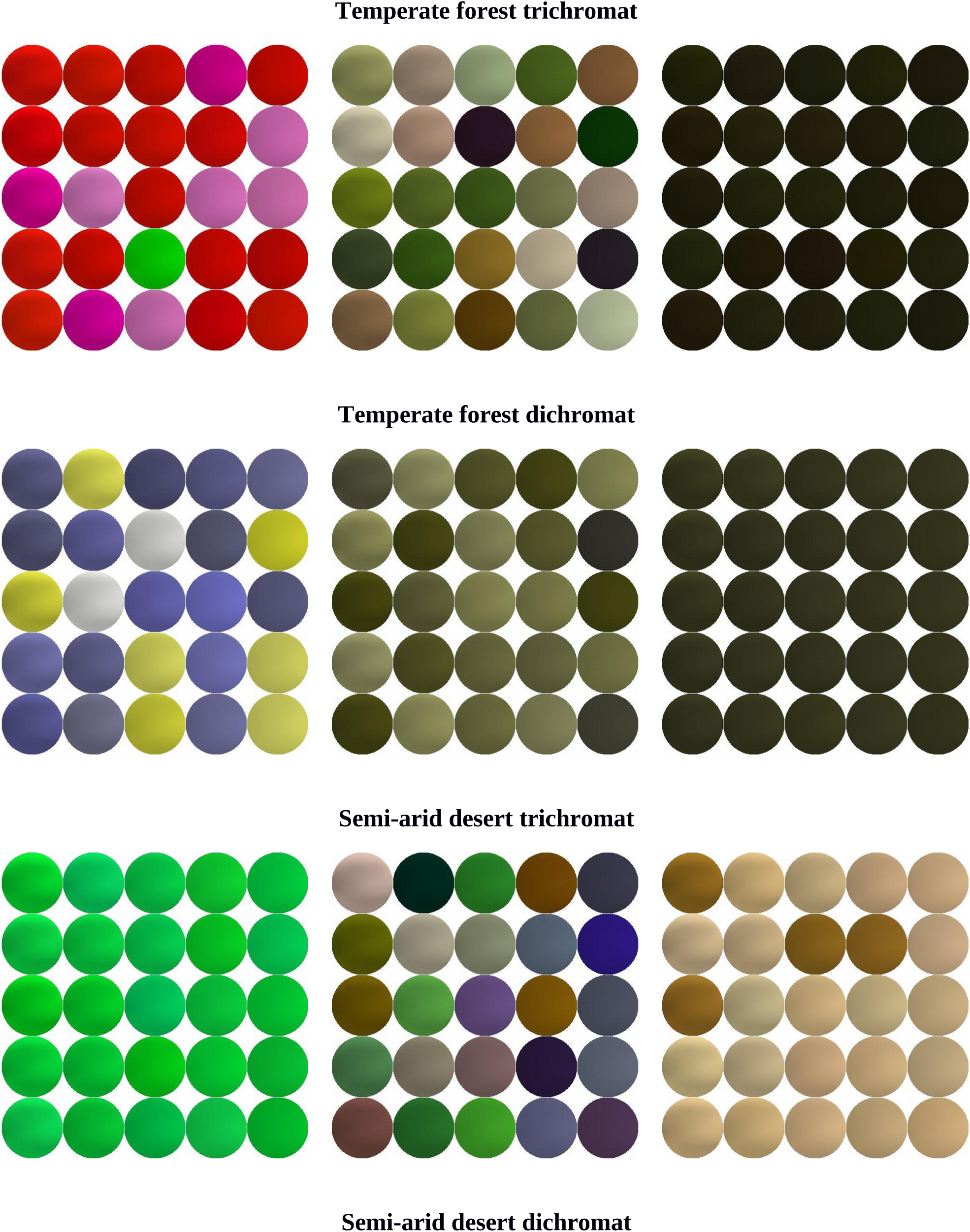
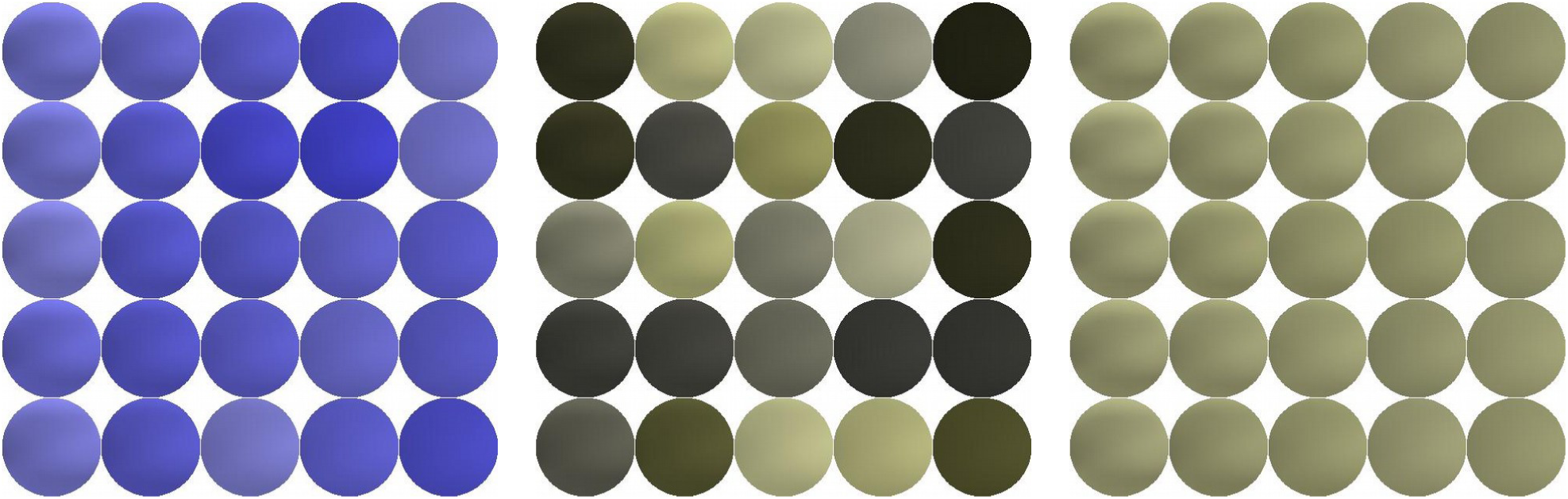
Representation of colours at similar reaction times. For each geographical location and colour condition the spheres represent, for illustration, 25 randomly sampled colours at a given reaction time (+/- 25 ms). The easiest to see colours are shown on the left, intermediate colours (chosen halfway between reaction time extremes) in the middle, and the hardest on the right. Reaction time steps were: temperate forest trichromat - 387 ms, 576 ms, 766 ms; temperate forest dichromat - 458 ms, 70 5ms, 952 ms; semi-arid desert trichromat - 374 ms, 485 ms, 595 ms; semi-arid desert dichromat - 429 ms, 605 ms, 781 ms.

## Discussion

Based on an approach using simple image synthesis, psychophysics and deep neural networks for interpolation, we identified the optimal colours for camouflage and conspicuity, an approach not previously tried for multidimensional perception-based experiments. We have shown that our method is capable of predicting entire parameter spaces and demonstrated its effectiveness with two and three dimensional colour spaces of considerable size. Furthermore, we provided confirmation that neural network predictions of hardest and easiest colours were consistent with human participants using a validation experiment. Interestingly, in some conditions the distribution with respect to hues appears to be multi-modal; for example, the hardest to find colours for the trichromat desert condition are either shades of dark beige or a lighter beige shade (resembling sand). This suggests that, unsurprisingly, there might be multiple solutions to the same problem, which intuitively seem to represent what is seen in natural and human-made camouflage [11]. In both the neural network predictions and the validation experiment, dichromatic targets were found to be significantly harder to detect than trichromatic targets for temperate forest and semi-arid desert conditions. Since our experiments were carried out using interleaved projected images of forest and desert scenes, it is important to rule out the possible confound of a greater switch cost between trials in different chromatic conditions (e.g. the switch cost is greater in the dichromat versus trichromat condition). To achieve this, we used an interstimulus interval of a mid-gray screen displayed for 2 seconds and checked that the mean luminance differences between trichromatic and simulated dichromatic images from the same geographical locations to the mean luminance of the interstimulus screen were not significantly different in both locations (temperate forest: *t*(62) = −0.5206, *p* = .6045; semi-arid desert: *t*(62) = −0.1397, *p* = .8893). Consequently this does not account for the difference between simulated dichromat and trichromat reaction times.

The result, that trichromatic vision is more effective at breaking camouflage, seems to run counter to oft-quoted historical accounts of the military value of dichromatic observers and contemporary theories for the maintenance of visual pigment polymorphisms in many New World monkey species [23,24]. However, the most recent work in this area suggests that evidence supporting an advantage for dichromats in camouflage breaking is, at best, equivocal [25]. This view seems to be confirmed by a brief survey of the literature. In one paper, Morgan, Adam and Mollon [26] review literature from as early as 1940 [27], which claims that dichromatic benefit would accrue in only limited situations, describing the early literature as largely descriptive and offering no empirical support. Morgan, Adam & Mollon [26] go on to describe their own empirical work with human observers, reporting dichromatic advantage, but their experiments were limited to a precisely controlled geometric display. Another study tested white-faced capuchins [28], arguing that some benefit accrues for dichromats, however, it is difficult to untangle the confounding effects of different light levels and relative abundance of target insects.

More ecologically relevant, Lovell et al. [29] investigated both trichromat and dichromat visual systems with respect to changes of illuminant in natural scenes, concluding that a foraging advantage accrues to trichromatic mammals because their visual system is less confounded by abrupt and unpredictable changes in illumination [p2069]; that is, it is less affected by shadows and changes in illumination. This is consistent with the present results.

An advantage for dichromats under particular conditions, but overall advantage for trichromats, seems to reflect the broad findings of this literature; indeed, it is the overall conclusion of Troscianko et al. [25]. They found that trichromats perform better, but under particular conditions dichromats have an advantage. Our own results arguably offer confirming evidence that, on average, across the whole colour gamut, trichromats perform better than dichromats, in two dissimilar environments. The intuitive statement below seems to sum up much of the empirical work that has been carried out:

“… *But for every instance of this kind that might be suggested, there are innumerable examples in which the colour-blind observer is at a marked disadvantage, and in other ways would of course be a source of real danger. Moreover, if the normal person were provided with pieces of coloured glass, it would be most unlikely that the colour-blind person would ever be able to score off him.*” [28]

Comparing the performance of trichromat and dichromat observers does not necessarily explain the visual ecology of real-world examples. What constitutes the best colour for camouflage of animals depends very much on the visual system of their prey and/or predator. Consider the coat of a tiger (*Felis tigris*); it has fur that appears orange to a trichromat observer rather than some shade of green, though the latter should be more appropriate camouflage for an ambush hunter in forests. However, as illustrated in the left-hand panel of Figure 4, when viewed as a dichromat, the tiger’s colour is very effective.

**Figure 4.**
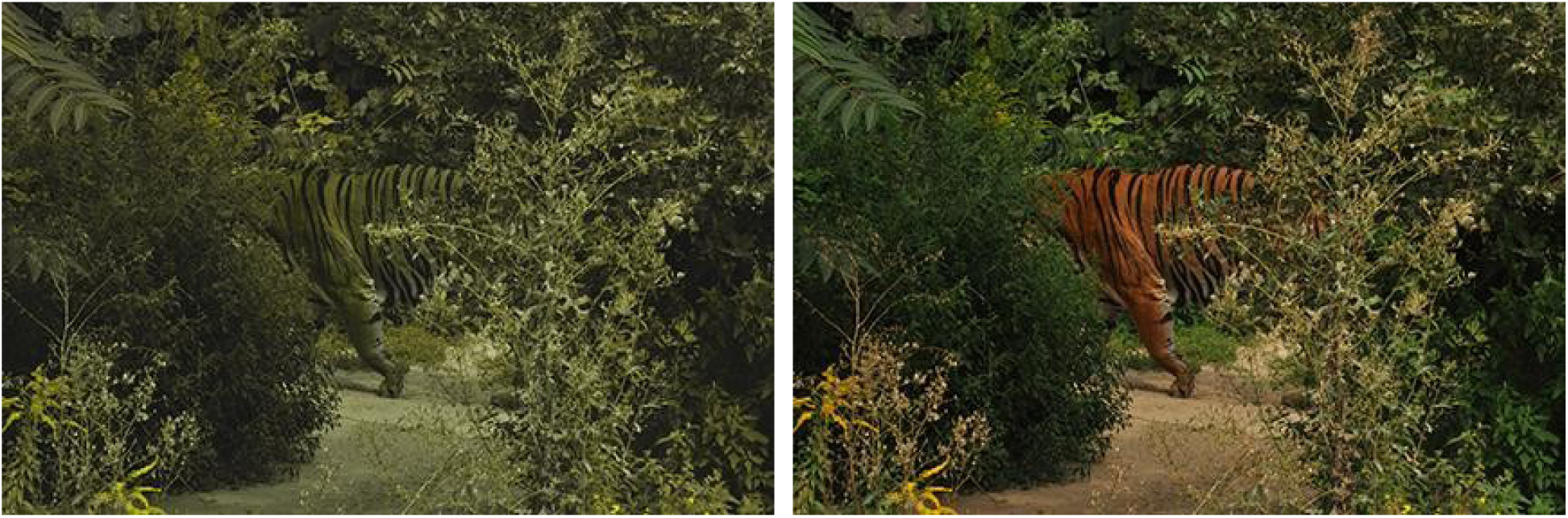
The effectiveness of the tigers colouring in the dichromat context is striking. Image of a tiger from the point of view of a dichromat receiver (left panel) and trichromat receiver (right panel).

## Conclusion

Based on our results and given that most non-human mammals have dichromatic colour vision that is unable to reliably differentiate orange and green, it seems that there is little benefit to actually become green if the receiver is dichromat. Hence predators (e.g. tigers), whose main prey is other mammals (e.g. deer), experience little evolutionary pressure to evolve green colouration from a trichromatic perspective. Deer are dichromats [30, 31] and, for them, most of their predators, like tigers, appear green. Moreover, producing a green coat would require a significant change to mammalian biochemistry since mammals rely on the large polymers, eumelanin and phaeomelanin, to produce black and yellow-red colours, which are the basis of the limited palette we see [32]. Indeed, the only mammal with a green coat is considered to be the sloth whose colour is actually due to a green alga (*Trichophilus welckeri*) that grows in its fur [33]. For species seeking concealment from dichromats there appears to be little pressure to actually become green. In contrast, when hiding from trichromats, simple colouration is not that effective. The open question is therefore not why predators are not green, but why their major prey are not trichromats.

## Method

### Participants

For collecting training data, five male and five female participants (each undertaking counterbalanced trichromat and dichromat experimental sessions) were recruited. Four participants (three female and one male) none of whom participated in the training data experiment took part in the validation experiment. All participants had normal or corrected-to-normal vision and were members of the University of Bristol. Informed consent was obtained from all participants as stated in the Declaration of Helsinki. All experiments were approved by the Ethics Committee of the University of Bristol’s Faculty of Science.

## Materials

### Stimulus construction (preliminaries)

Stimuli were created from three layers: (1) a foreground occlusion layer; (2) a target layer containing the search object; and (3) a background layer. (1) and (3) were taken from two locations, selected to represent two very different types of natural background (*temperate forest* in October 2015 in Leigh Woods, North Somerset, UK, 2° 38.6’ W, 51° 27.8’ N, and *semi-arid desert* in April 2016 in the Tabernas Desert, Almería, Spain, 2° 41.3’ E, 37° 02.9 N). Collection of the images for (3) consisted of choosing representative locations and taking 2848×4288 pixel photographs with a tripod mounted Nikon D90 digital SLR camera (Nikon Corp., Tokyo, Japan). Images for (1) were acquired using a large blue screen (1.8m x 2.8m blue cotton muslin photography background cloth mounted on a lightweight frame) that could be easily manoeuvred across the scene captured for (3). The blue-screen images were used with a chromakey technique to create occlusion of the search object during stimulus construction. Again, these images were captured using the tripod mounted Nikon D90 digital SLR.

The captured occlusion layer images were pre-processed in order to obtain the location of the blue screen as a mask. This provided for permissible locations of the centre of the search object that we wanted participants to find (see below), and identified all of those pixels that were needed to form the occlusion. This pre-processing allowed a location for the search object to be rapidly chosen and occlusion created during the experiment. The top panels of Figure 6 show the images used in pre-processing.

**Figure 6.**
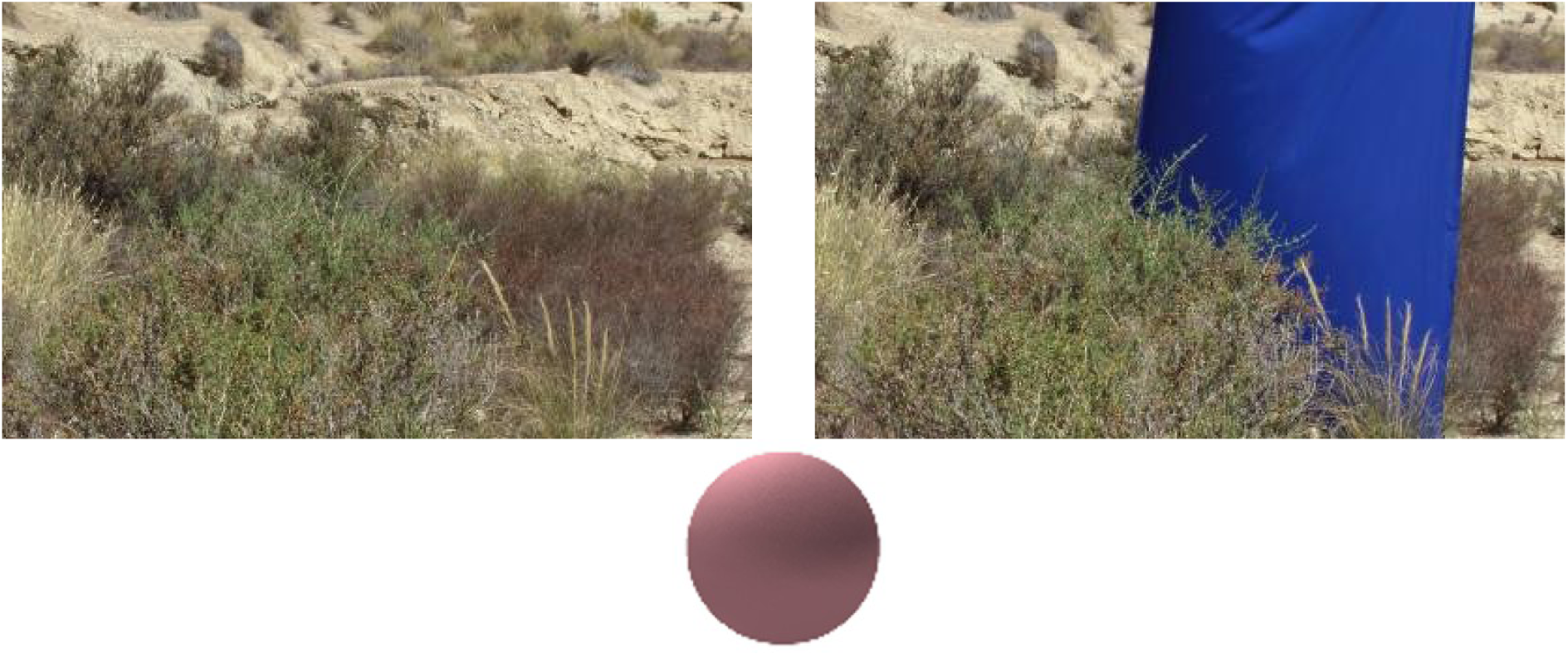
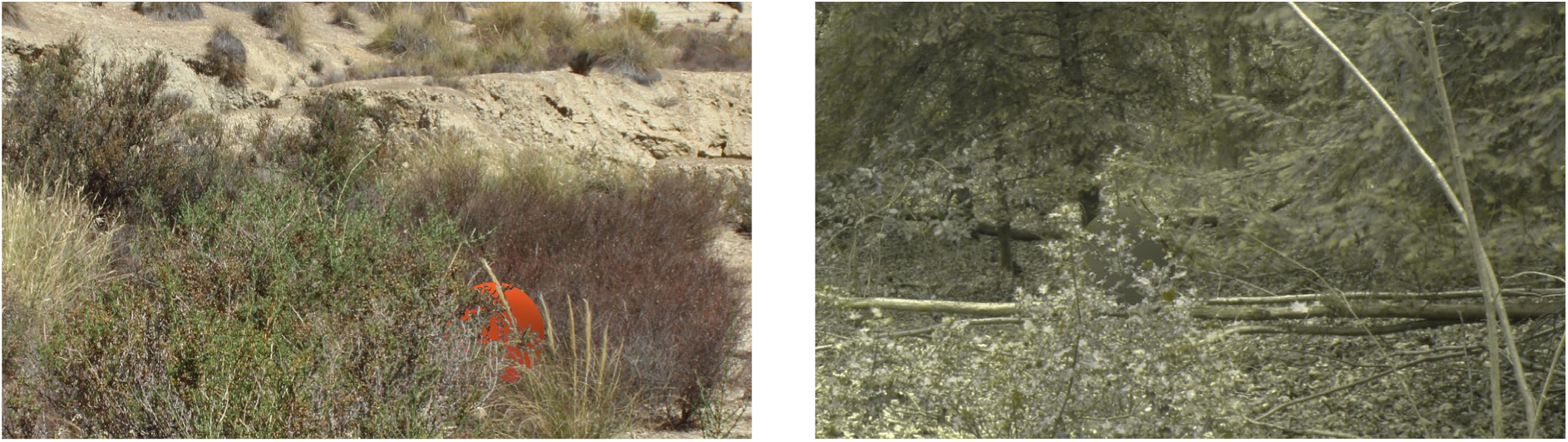
Examples of the background and blue screen images. Top left: a semi-arid desert background image; Top right: blue screen image using the same semi-arid desert scene; Centre: an example of a shaded sphere; Bottom left: an example of a trichromat stimulus displayed to participants using the semi-arid desert background images. Bottom right: a dichromat stimulus example using the temperate forest background images.

A bespoke program, written using Matlab 2017b (The MathWorks, Inc., Natick, MA, USA)and the Psychtoolbox-3 extensions [34,35], was used to construct and present the stimuli, and to collect experimental data.

### Stimulus construction (presentation)

During each trial, stimuli were constructed by randomly choosing a background image, together with its associated occlusion image, from a pool of 32 images and their permissible locations for the stimulus, for each geographical location. The occlusion image was pre-processed from the blue screen image and a matrix produced (the same size as the background image) containing a logical ‘true’ for every permissible centre for the search object. The combination of backgrounds (32 per location) and potential positions for the search image (mean 284,650 per image) provided a very large number of possible scenes. The target was a sphere, 128 pixels in diameter, constructed dynamically using a sample colour with pseudo-realistic shading to achieve a spherical look (an example of a coloured sphere is shown in the central panel of Figure 6). While we acknowledge that there are few perfectly round/spherical things in nature, we chose a sphere because it was easy to create and shade to look realistic. Maintaining a constant size and shape also had the benefit that any effects that might be attributable to changing shape could be discounted.

Based on the mask, a pixel position was randomly chosen as the centre point for the sphere and the sphere superimposed on the background. Also using the mask, the pixels occluding the sphere were then superimposed onto the sphere. Examples of the completed stimuli are shown in lower panels of Figure 6. Dichromatic representations of the stimuli were created using an implementation of the protan equation from Vienot, Brettel and Mollon [36], which creates a representation of a trichromat (RGB) image as perceived by people with protanopia.

### Procedure

Images were projected onto a 1900×1070mm screen (Euroscreen, Halmstad, Sweden) from 3100mm using a 1920×1080 pixel HD (contrast ratio 300,000:1) LCD projector (PT-AE7000U; Panasonic Corp., Kadoma, Japan). For Yxy measurements of projected colours, see Table S7 in Supplementary information. Participants sat behind a table 2 m from the display screen with a keyboard in front of them. The experimental stimulus subtended a visual angle of 60° by 33.75° and the target sphere 4°. Participants were randomly assigned to one of two colour space conditions which was presented in the first block (either trichromat or dichromat, the other condition being presented later on a separate occasion). A central fixation cross on a mid-grey background was displayed for 2 s prior to stimulus onset. Participants had up to 10 s to find and indicate on which side of the screen the stimulus sphere was presented. Failure to respond caused the trail to be recorded as a failure and the experiment to move on the next stimulus. Reaction times and errors were recorded. Each block consisted of 1000 trials (plus eight practice trials). Trials were based on 500 forest and 500 desert backgrounds presented in a random order. We used simple uniform random sampling without replacement to select sphere colours using a 24-bit RGB gamut. Occlusion levels were chosen randomly between 25 and 50%.

## Acknowledgements

LT and JGF were supported by an EPSRC Standard Grant no. EP/M006905/1 awarded to NESS, RJB and ICC. We thank Jasmina Stevanov for her help with the image acquisition in Spain and two anonymous referees for their helpful suggestions that improved the manuscript.

## Supplementary information

Details of network configuration, parameters and data analysis can be found in Supplementary information. Datasets for reaction times to stimuli are available online. Leigh Woods (temperate forest): **https://doi.org/10.6084/m9.figshare.7111214**. Tabernas Desert (semi-arid desert): **https://doi.org/10.6084/m9.figshare.7111217**.

