## Supplementary information for "Optimising colour for camouflage and visibility using deep learning: the effects of the environment and the observer’s visual system"

### Neural network architecture and parameters

Networks were trained for 500 epochs with a batch size of 128. The RMSprop optimiser was used with learning rate of 0.001 and mean squared error as loss function. The architecture of the networks is illustrated in Figure S1. All Dense layers had 768 units with ReLU activations and dropout was set to 0.5.

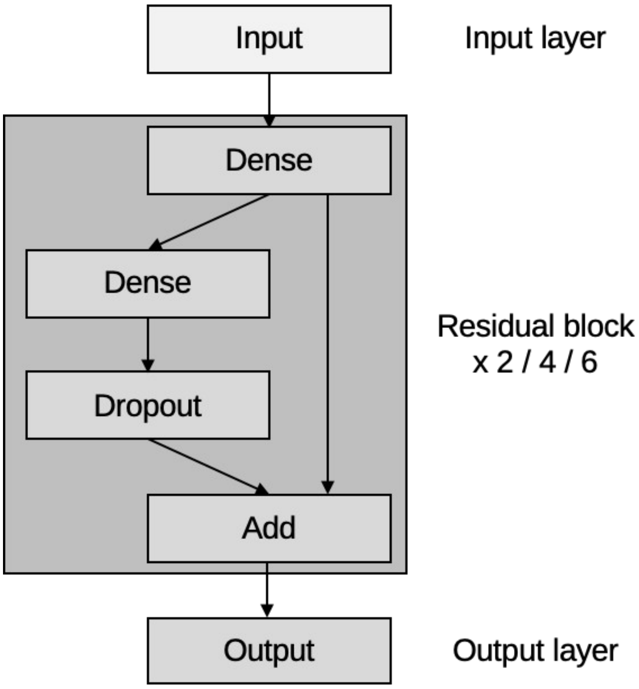


**Figure 1.** Schematic illustration of the residual deep neural networks used in the study.

### Comparing the error between networks with different number of residual blocks

Mean validation losses were calculated for 100 bootstrapped neural networks with two, four or six residual blocks after 500 training epochs using mean squared error (Fig. S2). Statistics were calculated using random permutation tests, based on 100,000 resamples. P-values were adjusted for multiple comparisons with False Discovery Rate [1]. We found that neural networks with four residual blocks produced significantly lower error rates compared to networks with two or six residual blocks, in all four experimental conditions (Table S1).

**Table S1.** Comparisons of mean validation losses for networks with two, four or six residual blocks in all four experimental conditions.

| Condition | Comparison | P-value |
|---|---|---|
| Temperate forest trichromat | 4 vs. 2 | .0093 |
| | 4 vs. 6 | .0093 |
| Temperate forest dichromat | 4 vs. 2 | .0012 |
| | 4 vs. 6 | .0115 |
| Semi-arid desert trichromat | 4 vs. 2 | < .0001 |
| | 4 vs. 6 | < .0001 |
| Semi-arid desert dichromat | 4 vs. 2 | .01022 |
| | 4 vs. 6 | .01022 |


2

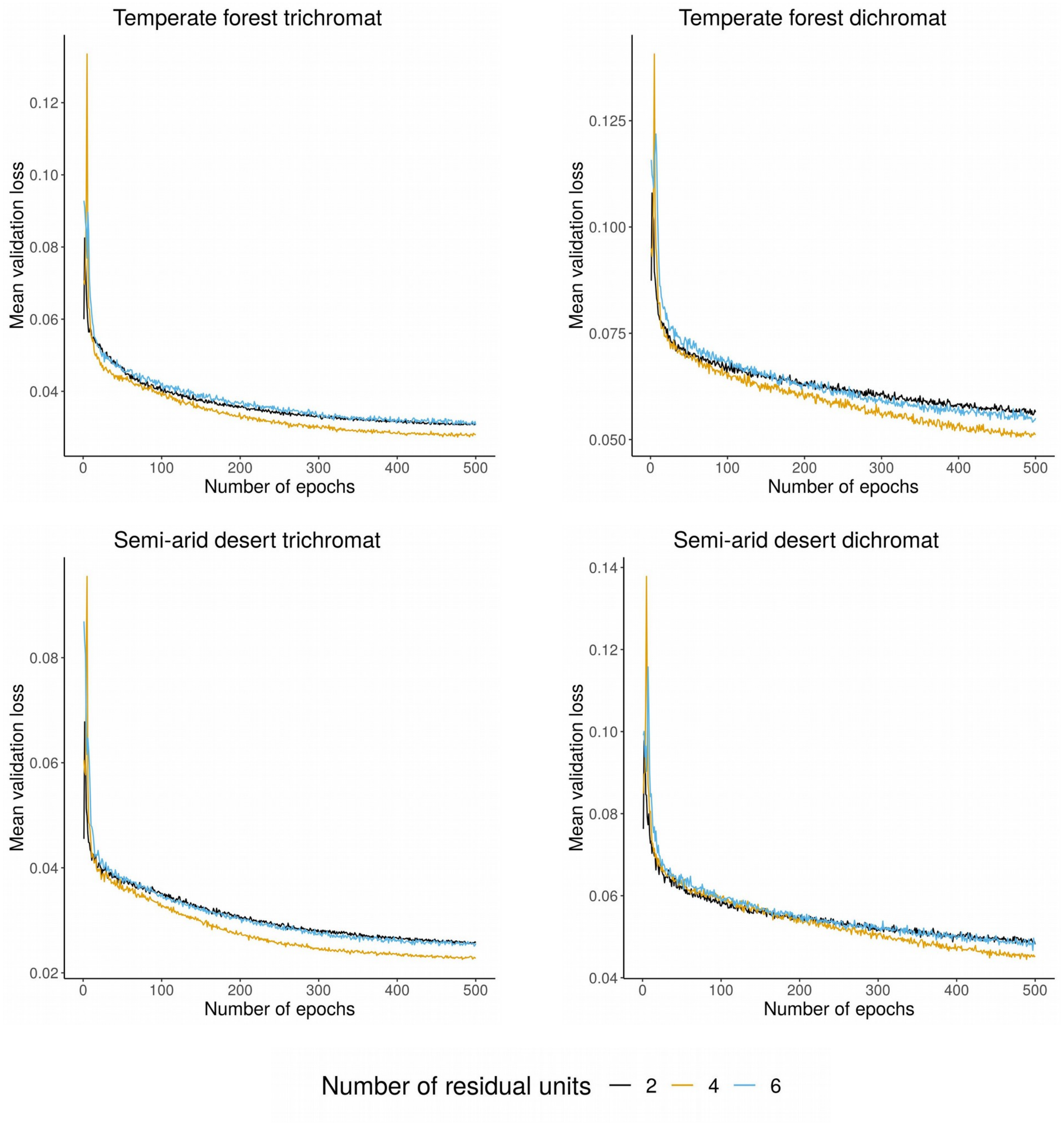


**Figure S2.** Mean validation losses for neural networks with two, four or six residual blocks across 500 training epochs for all four experimental conditions.

### GLMM results for trichromat vs. dichromat conditions in validation experiment

The effects of trichromat vs. dichromat conditions in the validation experiment were analysed by fitting generalised linear mixed models (GLMM) with gamma distributions (log link function) using the lme4 package [2] in R [3]. Gamma distributions were chosen due to the non-normality of reaction time data (Table S6) [4]. Nested models were compared using the change in deviance on removal of a term and by the Bayesian information criterion (BIC) [5]. Participant ID was treated as a random variable within the models. If a model including a term for chromatic condition had a significantly better fit to the one without it, the effect of the chromatic condition was significant (Table S2). Pairwise post-hoc analysis revealed that in all conditions dichromat targets were harder to see than trichromat targets (Table S3). P-values were adjusted for multiple comparisons with False Discovery Rate [1].

**Table S2.** Comparison of GLMMs with and without the chromatic condition term.

| Location | BIC (without) | BIC (with) | Δdeviance | DF | P-value |
|---|---|---|---|---|---|
| Temperate forest | -777.64 | -879.4 | 109.55 | 1 | < .0001 |
| Semi-arid desert | -2168.1 | -2204.7 | 44.443 | 1 | < .0001 |

**Table S3.** GLMM estimates, standard error and p-values from the post-hoc analysis of dichromat vs. trichromat conditions.

| Condition | Comparison | Estimate | Std. Error | P-value |
|---|---|---|---|---|
| Temperate forest | Dichromat Easiest vs. Trichromat Easiest | 0.0628 | 0.0248 | .0112 |
| | Dichromat Intermediate vs. Trichromat Intermediate | 0.1227 | 0.0248 | < .0001 |
| | Dichromat Hardest vs. Trichromat Hardest | 0.2726 | 0.0248 | < .0001 |
| Semi-arid desert | Dichromat Easiest vs. Trichromat Easiest | 0.0862 | 0.0213 | < .0001 |
| | Dichromat Intermediate vs. Trichromat Intermediate | 0.0627 | 0.0213 | .0032 |
| | Dichromat Hardest vs. Trichromat Hardest | 0.0898 | 0.0213 | < .0001 |

### GLMM results for increasing predicted difficulty in validation experiment

The effects of increasing predicted difficulty in the validation experiment were also analysed with GLMMs. If a model including a term for difficulty groups had a significantly better fit to the one without it, the effect of difficulty groups was significant (Table S4). Pairwise post-hoc analysis revealed that in all conditions progressively more difficult groups (predicted as easiest, intermediate, and hardest by the neural networks) were significantly harder to find (Table S5). P-values were adjusted for multiple comparisons with False Discovery Rate [1].

**Table S4.** Comparison of GLMMs with and without the difficulty groups term.

| Location | Chromatic condition | BIC (without) | BIC (with) | Δdeviance | DF | P-value |
|---|---|---|---|---|---|---|
| Temperate forest | Trichromat | -908.72 | -1068.74 | 174.20 | 2 | < .0001 |
| Temperate forest | Dichromat | 272.00 | 37.50 | 248.68 | 2 | < .0001 |
| Semi-arid desert | Trichromat | -1176.50 | -1283.00 | 120.68 | 2 | < .0001 |
| Semi-arid desert | Dichromat | -953.98 | -831.31 | 136.85 | 2 | < .0001 |

**Table S5.** GLMM estimates, standard error and p-values from the post-hoc analysis of difficulties in the validation experiment.

| Condition | Comparison | Estimate | Std. error | P-value |
|---|---|---|---|---|
| Temperate forest trichromat | Easiest vs. Intermediate | 0.2198 | 0.0203 | < .0001 |
| | Intermediate vs. Hardest | 0.0423 | 0.0203 | .0373 |
| Temperate forest dichromat | Easiest vs. Intermediate | 0.281 | 0.0284 | < .0001 |
| | Intermediate vs. Hardest | 0.1918 | 0.0284 | < .0001 |
| Semi-arid desert trichromat | Easiest vs. Intermediate | 0.0775 | 0.0194 | < .0001 |
| | Intermediate vs. Hardest | 0.1371 | 0.0194 | < .0001 |
| Semi-arid desert dichromat | Easiest vs. Intermediate | 0.1864 | 0.0206 | < .0001 |
| | Intermediate vs. Hardest | 0.0521 | 0.0206 | .0116 |

 5

**Table S6.** Shapiro-Wilk normality test results of the validation experiment.

| **Condition** | | **W** | **P-value** |
|---|---|---|---|
| Temperate forest trichromat | Easiest | 0.8186 | < .0001 |
| | Intermediate | 0.8235 | < .0001 |
| | Hardest | 0.866 | < .0001 |
| Temperate forest dichromat | Easiest | 0.8453 | < .0001 |
| | Intermediate | 0.864 | < .0001 |
| | Hardest | 0.7927 | < .0001 |
| Semi-arid trichromat | Easiest | 0.8623 | < .0001 |
| | Intermediate | 0.8418 | < .0001 |
| | Hardest | 0.7515 | < .0001 |
| Semi-arid dichromat | Easiest | 0.8620 | < .0001 |
| | Intermediate | 0.8785 | < .0001 |
| | Hardest | 0.8594 | < .0001 |

**Table S7.** Reference and measured values of projected colours using a Minolta CS-100A Luminance and Color Meter (Minolta Co., Ltd., Osaka, Japan).

| **Manufacturer's sRGB D65 colour values** | | | **Measured Yxy values** | | | **Description** |
|---|---|---|---|---|---|---|
| 52 | 53 | 53 | 4.0 | 0.278 | 0.333 | Black |
| 84 | 86 | 87 | 10.3 | 0.279 | 0.324 | Neutral 3.5 |
| 121 | 121 | 121 | 21.8 | 0.281 | 0.326 | Neutral 5 |
| 162 | 163 | 162 | 38.4 | 0.280 | 0.330 | Neaural 6.5 |
| 203 | 204 | 203 | 60.4 | 0.276 | 0.334 | Neutral 8 |
| 249 | 249 | 244 | 78.0 | 0.276 | 0.343 | White |
| 0 | 137 | 167 | 22.6 | 0.200 | 0.268 | Cyan |
| 190 | 87 | 152 | 17.2 | 0.294 | 0.202 | Magenta |
| 241 | 201 | 25 | 52.6 | 0.433 | 0.537 | Yellow |
| 174 | 60 | 61 | 10.6 | 0.501 | 0.326 | Red |
| 76 | 152 | 74 | 24.2 | 0.303 | 0.552 | Green |
| 49 | 68 | 151 | 8.3 | 0.167 | 0.138 | Blue |
| 230 | 160 | 45 | 27.4 | 0.465 | 0.497 | Orange yellow |


6

| | | | | | | |
|---|---|---|---|---|---|---|
| 162 | 190 | 65 | 34.5 | 0.380 | 0.563 | Yellow green |
| 93 | 61 | 105 | 6.2 | 0.250 | 0.201 | Purple |
| 195 | 83 | 97 | 13.2 | 0.423 | 0.296 | Moderate red |
| 72 | 92 | 174 | 11.0 | 0.178 | 0.159 | Purplish blue |
| 222 | 123 | 51 | 19.2 | 0.304 | 0.439 | Orange |
| 98 | 191 | 170 | 29.7 | 0.246 | 0.367 | Bluish green |
| 130 | 129 | 175 | 17.3 | 0.233 | 0.234 | Blue flower |
| 95 | 109 | 68 | 10.7 | 0.340 | 0.464 | Foliage |
| 92 | 123 | 156 | 16.2 | 0.220 | 0.253 | Blue sky |
| 195 | 147 | 129 | 23.8 | 0.356 | 0.354 | Light skin |
| 117 | 85 | 72 | 8.5 | 0.374 | 0.362 | Darkin skin |




7
